# Skeletal muscle cells derived from induced pluripotent stem cells: A platform for limb girdle muscular dystrophies

**DOI:** 10.1101/2022.05.08.489343

**Authors:** Céline Bruge, Marine Geoffroy, Manon Benabidès, Emilie Pellier, Evelyne Gicquel, Jamila Dhiab, Lucile Hoch, Isabelle Richard, Xavier Nissan

## Abstract

Limb girdle muscular dystrophies (LGMD), caused by mutations in 29 different genes, are the fourth most prevalent group of genetic muscle diseases, leading to progressive weakness and atrophy of the skeletal muscles. Although the link between LGMD and their genetic origins has been determined, LGMD still represent an unmet medical need. In this paper, we describe a platform for modeling LGMD based on the use of human induced pluripotent stem cells (hiPSC). Thanks to the self-renewing and pluripotency properties of hiPSC, this platform provides an alternative and renewable source of skeletal muscle cells (skMC) to primary, immortalized or overexpressing cells. We report that skMC derived from hiPSC express the majority of the genes and proteins causing LGMD. As a proof of concept, we demonstrate the importance of this cellular model for studying LGMDR9 by evaluating disease-specific phenotypes in skMC derived from hiPSC obtained from four patients.

## Introduction

Limb girdle muscular dystrophies (LGMD) refer to a heterogeneous group of rare genetic neuromuscular diseases, leading to progressive weakness and wasting of proximal pelvic and shoulder girdle muscles ^1–3^. Fourth most common group of muscular dystrophies with an estimated prevalence of 1.63 per 100,000 people ^4^, LGMD subtypes are caused by genetic alterations of various genes playing a critical role in muscle function, maintenance and repair ^5,6^. Since the first LGMD gene identification reported in 1995 ^7^, the number LGMD subtypes has grown to a total of 29 according to the last revised nomenclature ^8^. Classified according their inheritance pattern, LGMD subtypes are autosomal dominant (LGMD D) or autosomal recessive (LGMD R) types affecting 27 different proteins of the nucleus, sarcoplasm, sarcomere, sarcolemma, and extracellular matrix (Table S1). While several gene replacement therapies ^9–14^ have been proposed over the years, no curative treatment is currently available for any types of LGMD.

Over the past decade, considerable efforts have been reported for deciphering the molecular mechanisms associated with LGMD. Understanding the multiple molecular pathways associated with LGMD requires the development of appropriate cellular and animal models that at least partially recapitulate the characteristics of these diseases. Animal models are currently preferred to *in vitro* models for studies on muscular dystrophies ^15,16^ and the assessment of therapeutic strategies ^17^. However, although most of these animal models recapitulate the pathological phenotypes observed in humans, some of these models appear relatively normal or with a mild dystrophic phenotype ^18^, or even present impairments that are not reported in patients ^19^. Alternatively, over the past decades, several *in vitro* human models have been developed to study phenotypes associated with LGMD. Among these, primary skeletal muscle cells (skMC) from muscle biopsies represent the most commonly reported biological resource, despite the difficulty of their generation and their limited proliferative capacity ^20^. To overcome these issues, several *in vitro* models were developed through transdifferentiation of non-muscle cells into skeletal cells by forced expression of myogenic transcription factors ^21^. More recently, immortalization of primary cells with hTERT and CDK4 was reported as successful for modeling LGMD ^22,23^, including LGMDD4/R1 ^24^, LGMDR2 ^25–27^ and LGMDR12 ^28^ While these different models present several advantages in terms of proliferation and maturity, the development of transgene-free cellular models from non-invasive procedures was proposed through the use of pluripotent stem cells.

Derived from embryos ^29–32^ or generated through reprogramming ^33,34^, pluripotent stem cells have the unique capacity to unlimitedly self-renew and differentiate into any cell type in the organism ^35^. Over the years, thanks to the development of efficient protocols of differentiation for a growing number of cell types, embryonic stem cells (ES) and human induced pluripotent stem cells (hiPSC) have been reported as efficient alternative models to primary or genetically modified cells. Since their discovery in humans, in 1998 and 2007, the use of these cells has been explored for a broad range of applications, leading to the identification of new molecular mechanisms ^36,37^, new pathological phenotypes ^38–41^ and the development of new therapies ^42–45^ (reviewed in Karagiannis et al., 2019) ^46^. Although successfully applied for a number of genetic diseases, pathological modeling using pluripotent stem cells remained challenging for application to muscular disorders over almost a decade because of the lack of protocols for generating homogeneous populations of myotubes. Since 2012, several studies have reported methods to overcome this limitation by differentiating hiPSC into myogenic cells using transgene-based methods ^47–49^ or small molecules recapitulating developmental myogenesis ^50–57^. Among these protocols, Caron et al. described a three-step process that efficiently differentiates hiPSC into mature skMC in less than 26 days using combinations of small molecules and growth factors identified by high-throughput screening. Recently, several studies have highlighted the use of human pluripotent stem cells, and more particularly hiPSC, for studying neuromuscular disorders ^58–62^, including LGMD ^49,63–66^. In the present study, we report that hiPSC-derived skMC provide a reliable and robust *in vitro* tool for investigating LGMD subtypes. To do so, we used a robust protocol of differentiation to generate pure populations of skeletal myoblasts (skMb) and terminally differentiated myotubes (skMt) from hiPSC ^51^. Taking advantage of this model, we monitored and characterized the expression and localization of all the genes and proteins causing LGMD subtypes. As a proof of concept, such a cellular model was used to model LGMDR9 by analyzing alpha-dystroglycan (α-DG) glycosylation in fukutin-related protein (FKRP)-deficient skMt derived from the hiPSC of four patients.

## Results

### Generation and characterization of skeletal muscle cells from human induced-pluripotent stem cells

The workflow to generate skeletal myoblasts cells (skMb) and multinucleated myotubes (skMt) from hiPSC is represented in Figure 1A. First, pluripotency capacities of three healthy hiPSC lines (WT1,2,3) were characterized by analyzing the expression of Oct4 and Nanog through immunostaining (Figure 1B) and quantifying SSEA4/TRA1-80 by flow cytometry (Figure S1). Myogenic differentiation was then induced using a 3-step protocol of differentiation ^51^. Briefly, SKM01 medium was used for 10 days to initiate the differentiation of hiPSC into myogenic precursors before their maturation into myoblasts in SKM02 medium and their terminal differentiation into myotubes in SKM03 medium. To demonstrate the efficacy and robustness of this protocol of differentiation, MyoD and desmin expressions were assessed by immunostaining in skMb, showing homogeneity of labeling (Figure 1C). Terminal differentiation of skMt was then characterized by measuring the expression of myogenin and titin by immunostaining (Figure 1D). To quantify gene expression changes throughout the differentiation protocol, the same markers were monitored by qPCR, revealing, on the one hand, a significant time dependent decrease of Oct4 and Nanog in skMb and skMt compared to hiPSC (Figure 1E) and, on the other hand, a significant induction of MyoD, desmin, MyoG and titin in skMb and skMt (Figures 1F and 1G).

**Figure 1.**
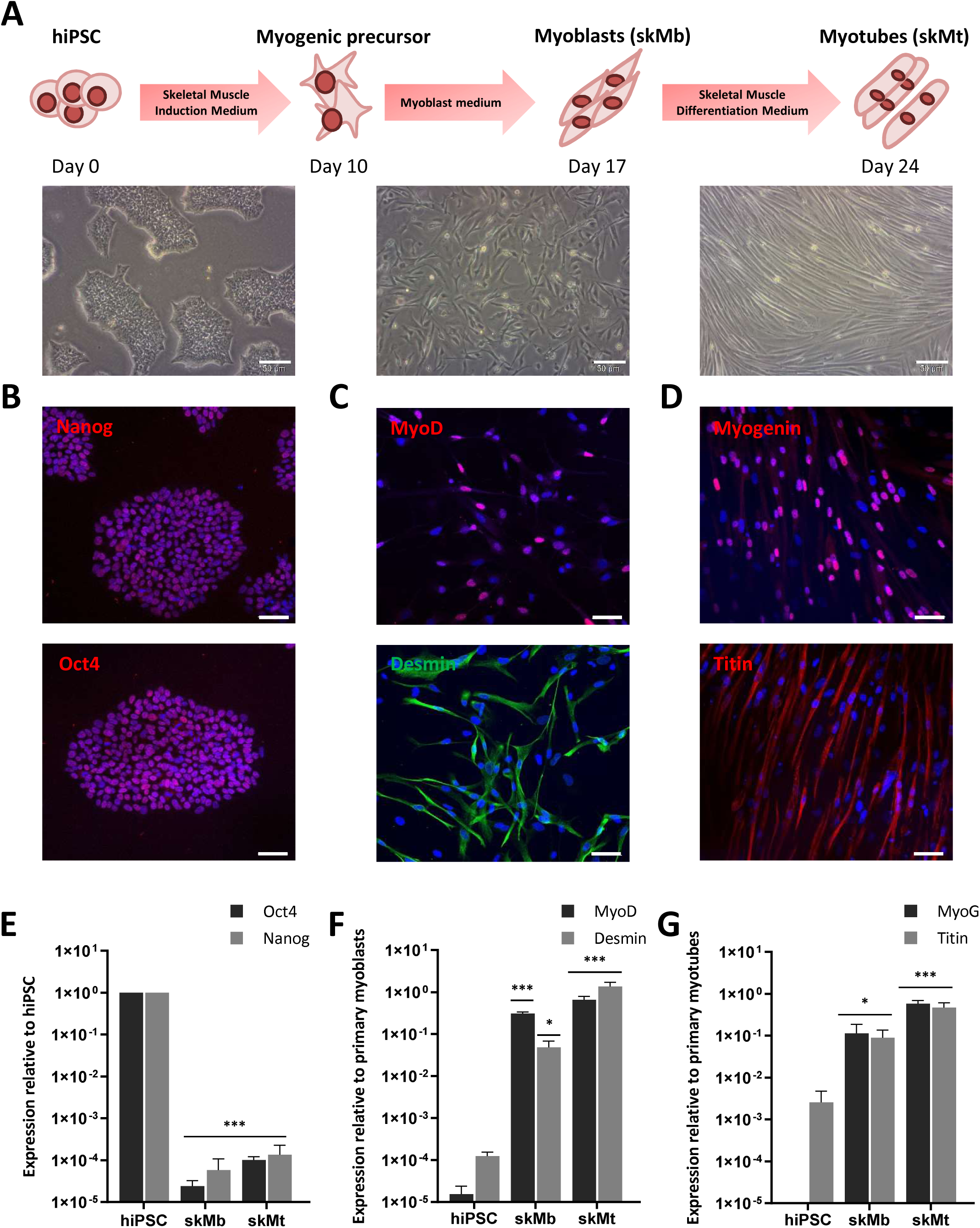
Differentiation of human induced pluripotent stem cells (hiPSC) into skeletal muscle cells (skMC). **(A)** Experimental workflow of skMC differentiation from hiPSC using the Genea Biocells protocol. Representative pictures of hiPSC, skeletal myoblasts (skMb) and myotubes (skMt) with bright field acquisition. Scale bar = 50 µm. **(B)** Characterization of the expression of pluripotency markers Oct4 and Nanog by immunostaining in hiPSC. **(C)** Characterization of the expression of MyoD and desmin by immunostaining in skMb **(D)** Characterization of the expression of MyoG and titin by immunostaining in skMt. Nuclei are labelled by Hoechst staining (blue). Scale bar = 50 µm. **(E)** Measurement of Oct4 and Nanog expression by qPCR in hiPSC, skMb and skMt. **(F)** Measurement of MyoD and desmin expression by qPCR in hiPSC, skMb and skMt. **(G)** Measurement of MyoG and titin expression by qPCR in hiPSC, skMb and skMt. Gene expression analyses are normalized to hiPSC, primary myoblasts or primary myotubes. Data are shown as the mean of skMC differentiations of three control cell lines +/- SD. ***p≤0.005 (Student’s t-test). Abbreviations: hiPSC, human induced pluripotent stem cells; skMb, skeletal myoblasts; skMt, skeletal myotubes; MyoG, myogenin.

To further characterize the terminal differentiation of skMt derived from hiPSC and visualize sarcomeric structure formation in real time, we then used two gene-edited hiPSC lines expressing titin (TTN) or troponin I (TNN) proteins fused to GFP (AICS-48 and AIC-37). After having validated their pluripotency capacities by measuring the expression of SSEA4/TRA1-80 using flow cytometry (Figure S2A), TTN-GFP and TNN-GFP hiPSC were differentiated using the same protocol. Analysis by qPCR confirmed the efficient myogenic differentiation of these cells by showing a significant decrease of Oct4 and Nanog expression in skMb and skMt compared to undifferentiated hiPSC (Figure S2B) and a significant increase of MyoD, desmin, MyoG and titin in skMb and skMt (Figure S2B). Terminal differentiation of TTN-GFP and TNN-GFP skMt was then followed in real time using a time-lapse analysis of GFP expression from Day 0 to Day 7 (Movie S1 and S2). Confocal microscopy analysis of GFP expression and localization in skMt derived from these two hiPSC cell lines revealed the appearance of GFP in both cell lines from Day 3 onwards (Figure 2A). Quantification of GFP positive cells showed that half of the cells expressed GFP from Day 3 of maturation and up to 80% at Day 5 (Figures 2B and 2C). Organization of the sarcomeric structures in skMt was further analyzed by characterizing the Z-disk, I-band and M-line using specific antibodies, as shown in Figure 2D. Confocal analysis of immunostaining and the GFP signal was performed in TTN-GFP skMt (Figure 2E) and TNN-GFP skMt (Figure S2C), revealing a regular, transverse and repetitive organization of sarcomeres. Finally, quantification of TTN-GFP skMt presenting a striated pattern revealed a time dependent maturation from 40% at Day 2 to 80% at Day 5 (Figure 2F).

**Figure 2.**
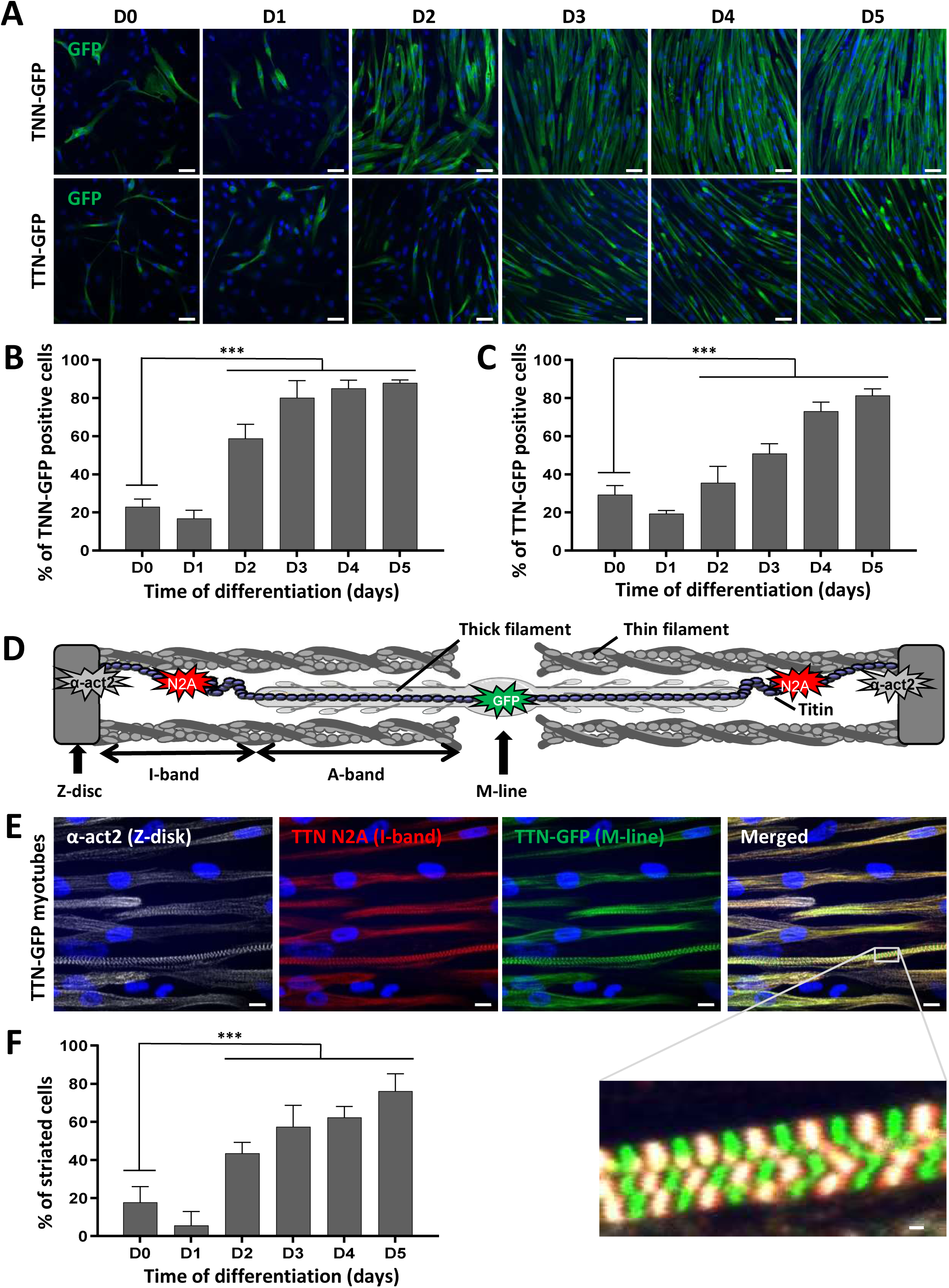
Striation pattern appears during terminal myogenic differentiation. **(A-C):** Troponin I(TNN)- and titin(TTN)-eGFP dynamics during myotube maturation. **(A)** Confocal images of TNN- and TTN-eGFP tagged cell lines were obtained from Day 0 to Day 5 after induction of myotube maturation. Nuclei are labelled by Hoechst staining. Scale bar = 50 µm. Quantification of the percentage of cells terminally differentiated, presented as the percentage of **(B)** TNN- or **(C)** TTN-positive cells out of the total number of myotubes. Each chart represents the mean +/- SD (n=5) for a representative experiment; ***p≤0.005 (Student’s t-test). **(D-F):** Analysis of sarcomeric organization during myotube maturation. **(D)** Schematic representation of sarcomeric architecture in skeletal muscle: Z-disk composed of α-actinin, I-band made up of actin thin filaments, A-band comprised of myosin thick filaments and the M-line where filaments are anchored. Titin spans the entire half-sarcomere: from the Z-disk with the N-terminal extremity to the M-line, by way of the I-band specifically recognized by the titin N2A antibody. **(E)** Immunofluorescent staining and confocal acquisition of sarcomere structures with TTN-eGFP-tagged cells at Day 5 of myotube maturation. Nuclei are labelled by Hoechst staining. Scale bar = 10 µm. White box denotes an enlargement of co-staining to visualize the cross-striation pattern of myotubes. Scale bar = 1 µm. **(F)** Quantification of the number of striated cells relative to the total number of skMt during time course differentiation of the TTN-eGFP-tagged cell line. Each chart represents the mean +/- SD (n=10) for a representative experiment; ***p≤0.005 (Student’s t-test). Abbreviations: TNN, troponin I; TTN, titin; α-act2, alpha-actinin2.

### Characterization of LGMD genes and proteins in skMt derived from hiPSC

Mutations in 27 genes are reported to cause 29 distinct LGMD (Table 1). Expression of these 27 genes was measured by qPCR in skMt derived from three healthy hiPSC lines (WT1,2,3) and myotubes derived from four healthy immortalized myoblast cell lines (C25, AB678, AB1079, AB1167) (Figure 3). Given that collagen type 6 is composed of three genetic components, alpha 1 to alpha 3 chains, three genes were monitored for this LGMD subtype, resulting in a total of 29 genes measured by gene expression in these two sources of myotubes. Analysis of this qPCR revealed similar levels of expression for the majority of these genes, both in skMt and immortalized myotubes: DNAJB6, TNPO3, HNRNPDL, CAPN3, COL6A3, DYSF, SGCB, SGCG, SGCD, TCAP, TRIM32, POMT1, FKTN, POMT2, DAG1, TRAPPC11, GMPPB and POGLUT1. Among these 29 genes, 2 genes revealed significant differences (10x fold change): on the one hand, *ANO5*, that was upregulated in skMt and, on the other hand, *LAMA2*, that was significantly upregulated in immortalized myotubes.

**Figure 3.**
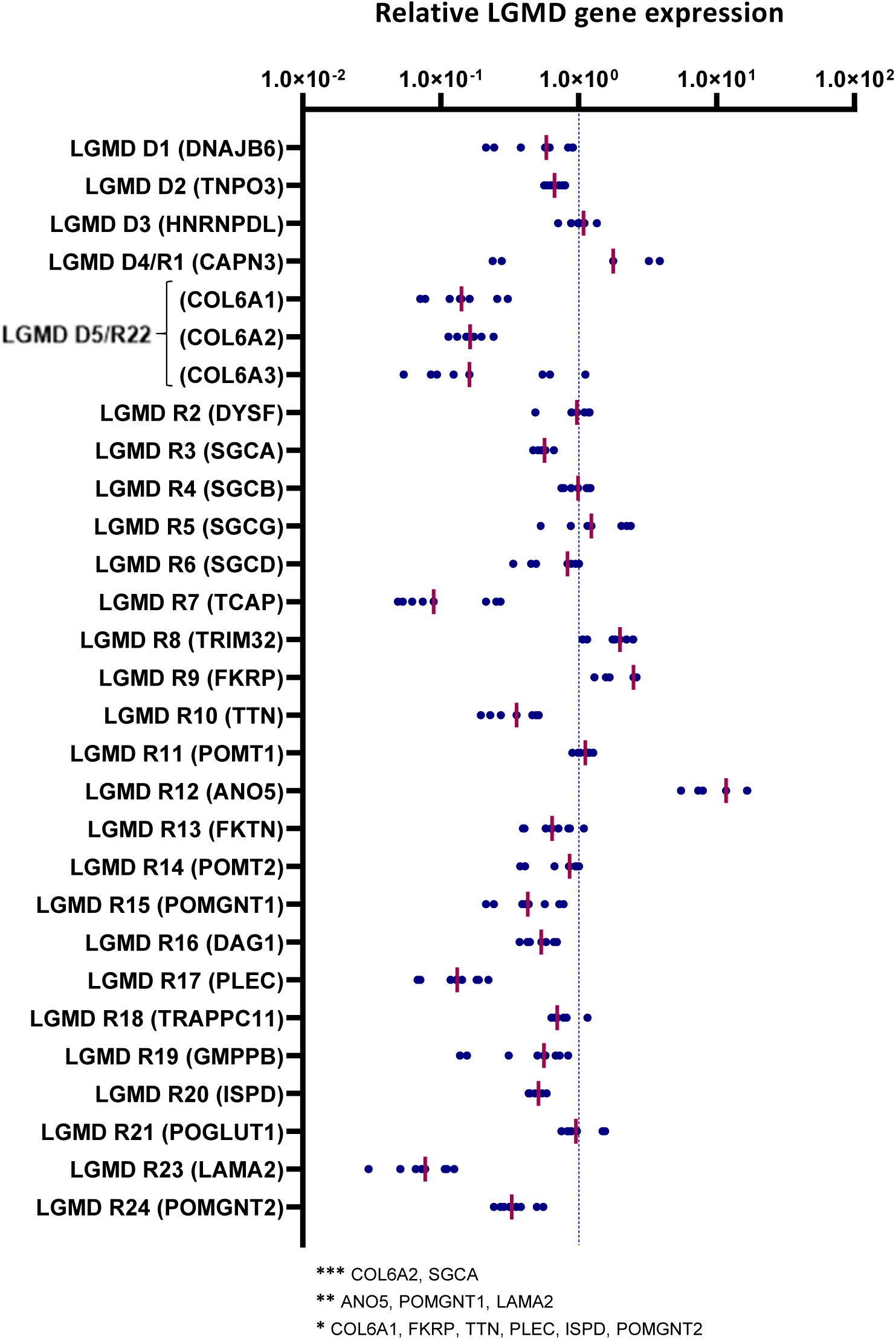
Gene expression analysis of LGMD genes in skMt. Gene expression analysis of LGMD genes in skMt derived from three healthy hiPSC lines (WT1,2,3) at Day 7 of myotube maturation. Fold changes (on log 10 scale) for ΔΔCT, relative to myotubes derived from 4 healthy primary myoblast cell lines (C25, AB678, AB1079, AB1167), are plotted as a group of data points and organized by gene according to the last revised LGMD classification. The pink horizontal line in each group of data points represents the mean for the group; *0.01<p≤0.05 or **0.005<p ≤ 0.01 or ***p<0.005. Abbreviations: ANO5, anoctamin 5; CAPN3, calpain 3; COL6, collagen 6; DAG1, dystroglycan 1; DNAJB6, DnaJ heat shock protein family member B6; DYSF, dysferlin; FKRP, fukutin-related protein; FKTN, fukutin; GMPPB, GDP-mannose pyrophosphorylase B; HNRNPDL, heterogeneous nuclear ribonucleoprotein D-like; ISPD, isoprenoid synthase domain-containing protein; LAMA2, laminin subunit alpha 2; PLEC, plectin; POGLUT1, protein O-glucosyltransferase 1; POMGNT, protein O-linked mannose N-acetylglucosaminyltransferase; POMT, protein O-mannosyltransferase; SGC, sarcoglycan; TCAP, titin-cap; TNPO3, transportin 3; TRAPPC11, trafficking protein particle complex subunit 11; TRIM32, tripartite motif containing 32; TTN, titin.

We then evaluated the expression of the corresponding proteins by immunostaining in skMt derived from one representative control hiPSC line (Figure 4). Confocal analysis of the expression of the dystrophin-associated glycoprotein complex (DGC) components revealed positive staining of dystroglycan 1 (DAG1) and beta, delta, gamma sarcoglycans (SGCB, SGCD and SGCG), while the alpha subunit (SGCA) was not detected in skMt. Analysis of sarcomeric components showed positive expression and localization within sarcomeres of calpain-3 (CAPN3), telethonin (TCAP) and titin (TTN), while a more diffuse, cross-striated pattern of plectin (PLEC) was also detected in skMt. We then evaluated the expression of proteins involved in DG glycosylation. Immunostaining of POMT2, POGLUT1 and FKRP showed a punctiform perinuclear localization of proteins, while localization was cytoplasmic for POMGnt1 and 2, POMT1, FKTN, GMPPB and ISPD. Regarding the proteins involved in trafficking, analysis of dysferlin expression by immunostaining revealed the presence of the protein at the plasma membrane and also in the cytoplasm of skMt, whereas TNPO3 and TRAPPC11 labeling was cytoplasmic. Staining of DNAJB6 resulted in a signal in the nucleus and the cytoplasm, whereas that of anoctamin 5 (ANO5/TMEM16E) and tripartite motif-containing protein 32 (TRIM32) revealed a punctiform signal in the cytoplasm. With regard to the nuclear HNRNPDL protein, staining was observed in the nucleus of skMt, as expected. Finally, analysis of the extracellular matrix proteins performed in non-permeabilized condition confirmed the presence of laminin-2 (LAMA2) in the sarcolemma. Immunostaining of collagen-6 alpha subunits 1, 2 and 3 (COL6A1, COL6A2, COL6A3) revealed a punctiform signal across skMt. Altogether, qPCR and immunostaining data show that hiPSC-derived skMt express the majority of LGMD genes at a comparable level to immortalized myotubes and that the related encoded proteins are present in the expected cellular compartment.

**Figure 4.**
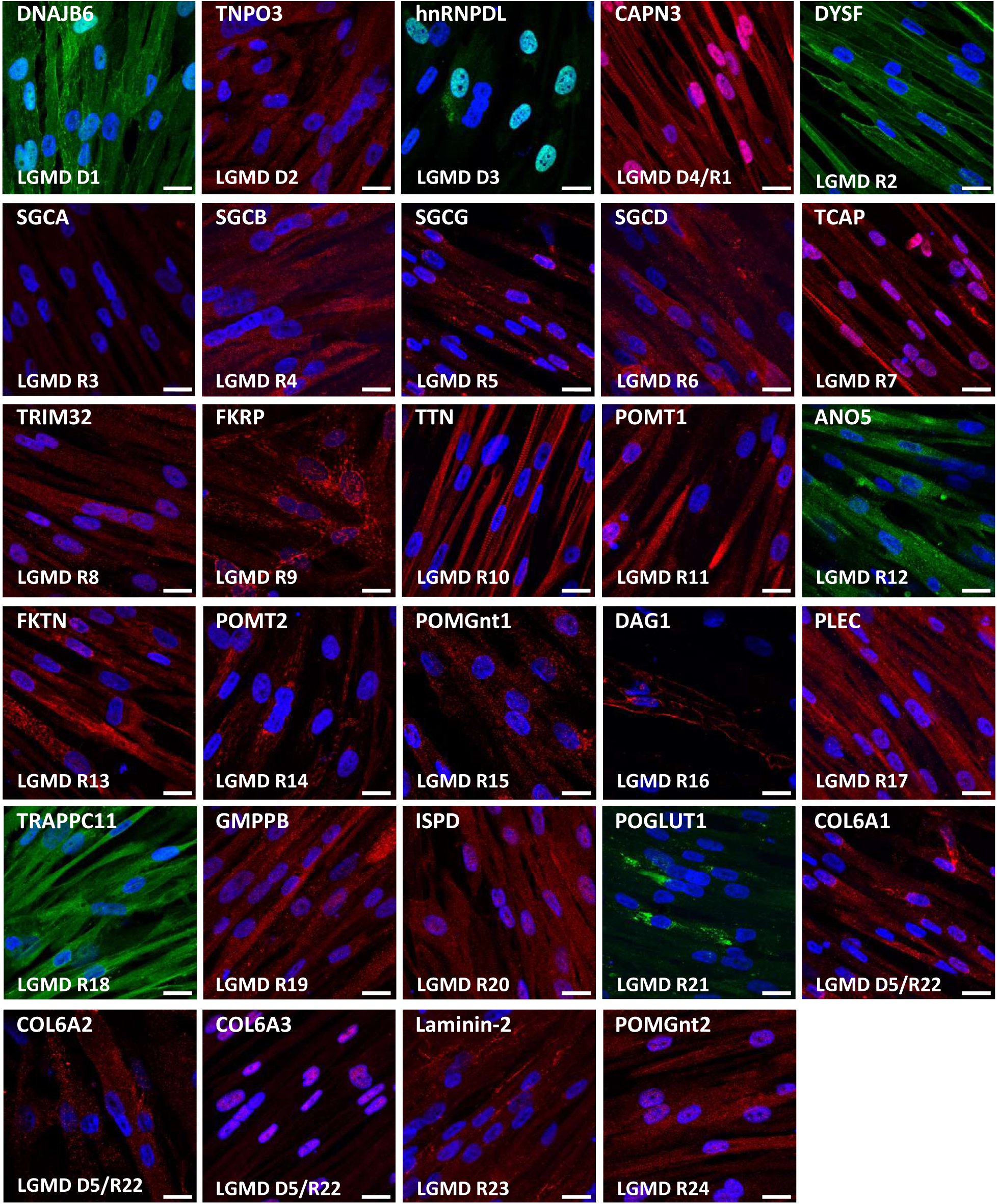
Characterization of LGMD proteins in skMt. Representative images of LGMD protein immunostaining in skMt derived from one control hiPSC line at Day 5 of maturation. Nuclei are labelled by Hoechst staining. Scale bar = 20µm. Abbreviations: ANO5, anoctamin 5; CAPN3, calpain 3; COL6, collagen 6; DAG1, dystroglycan 1; DNAJB6, DnaJ heat shock protein family member B6; DYSF, dysferlin; FKRP, fukutin-related protein; FKTN, fukutin; GMPPB, GDP-mannose pyrophosphorylase B; hnRNPDL, heterogeneous nuclear ribonucleoprotein D-like; ISPD, isoprenoid synthase domain-containing protein; LAMA2, laminin subunit alpha 2; PLEC, plectin; POGLUT1, protein O-glucosyltransferase 1; POMGnt, protein O-linked mannose N-acetylglucosaminyltransferase; POMT, protein O-mannosyltransferase; SGC, sarcoglycan; TCAP, titin-cap; TNPO3, transportin 3; TRAPPC11, trafficking protein particle complex subunit 11; TRIM32, tripartite motif containing 32; TTN, titin.

### Derivation and characterization of skMt from FKRP-deficient hiPSC

As a proof of concept, we then investigated the pathological features of LGMDR9 using hiPSC. To do so, an hiPSC cell line from one patient was provided by the LGMD2I Research Fund and three additional hiPSC lines carrying mutations in the *FKRP* gene were generated by reprogramming patient cells. Pluripotency of the four LGMDR9 hiPSC lines was confirmed by measuring the expression of Oct4 and Nanog using qPCR (Figure S3A) and immunostaining (Figure 5A). Pluripotency status of these hiPSC lines was also validated by measuring the expression of SSEA4 and TRA1-80 using flow cytometry (Figure S3B). Finally, chromosomal stability was analyzed using the multiplex fluorescence in situ hybridization (m-FISH) technique, revealing the absence of significant abnormalities (Figure S3C). Following this quality control assessment, the four FKRP-deficient hiPSC lines were further differentiated into skMb and skMt and compared to one representative healthy hiPSC line. Molecular characterization of skMb derived from FKRP-deficient hiPSC revealed no difference from the healthy skMb in terms of MyoD expression (Figure S4A). FKRP-deficient skMb were secondarily differentiated into skMt, showing positive sarcomeric staining for myosin heavy chain (MHC), titin and α-actinin (Figures 5B and S4B) suggesting that the absence of FKRP does not affect either the initial or the terminal stages of differentiation. Analysis of α-DG glycosylation was then performed by immunostaining in FKRP-deficient skMt and compared to the representative healthy relatives. Confocal analyses of IIH6 staining revealed the absence of a glycosylated α-DG signal in the FKRP1 line (V300A/A321E mutation), but not in the other cell lines (FKRP2,3,4), when compared to the healthy control (Figure 5B). This result was subsequently confirmed by western blot, showing that IIH6 staining was decreased in the same cell line (Figure 5C).

**Figure 5.**
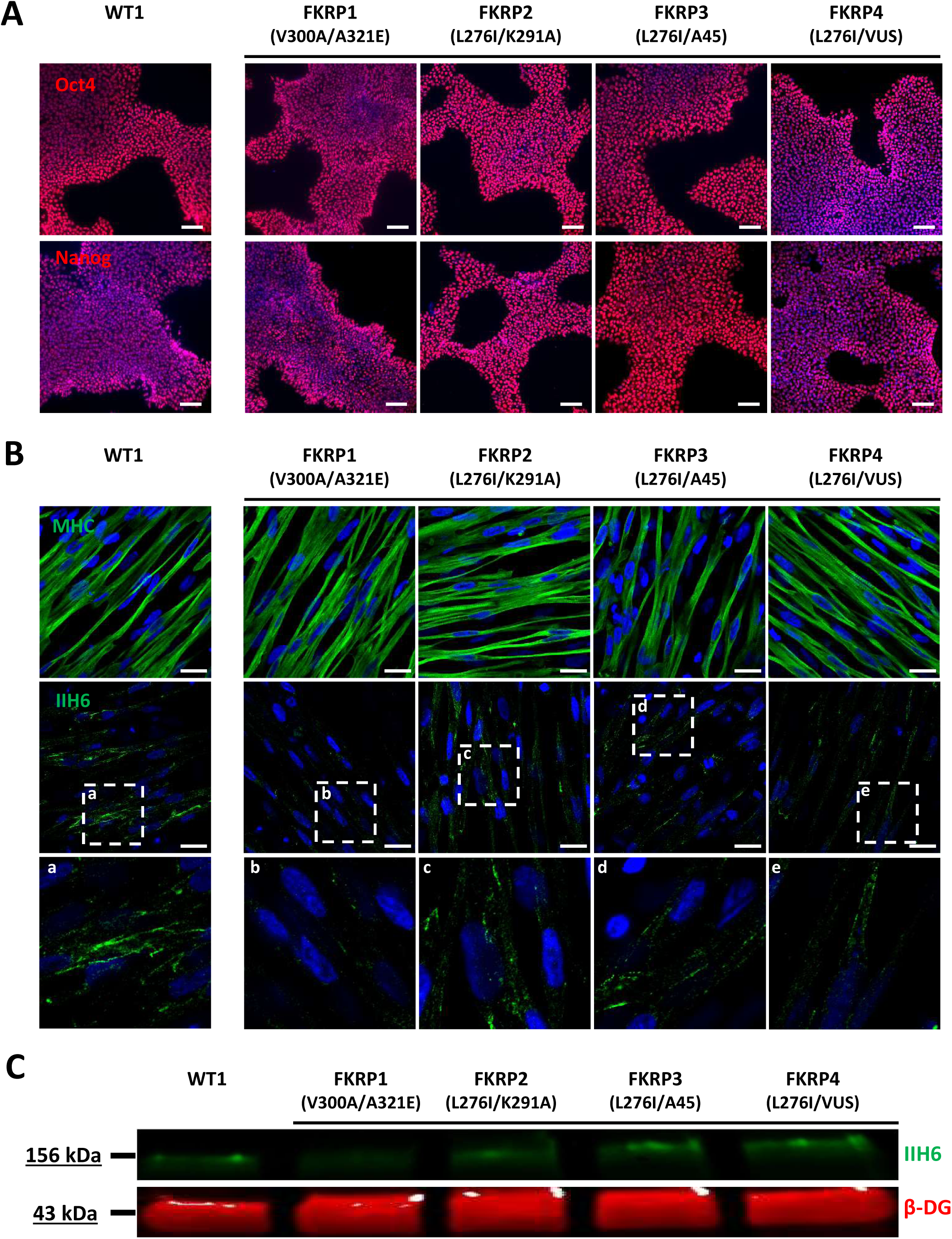
Differentiation and characterization of LGMDR9-affected hiPSC into SkMC. **(A)** Immunostaining analysis of Oct4 and Nanog in one control and four LGMDR9 hiPSC lines at Day 0 of differentiation. **(B)** Immunostaining images of MHC (upper panel) and α-DG glycosylation with IIH6 antibody (middle panel) in one control and four LGMDR9-derived skMt at Day 24 of differentiation. White boxes delineate enlargement of IIH6 staining to visualize higher magnification of α-DG glycosylation in the bottom panel. Nuclei are labelled by Hoechst staining. Scale bar = 20 µm. **(C)** Immunoblot analysis of α-DG glycosylation in skMt at Day 24 of differentiation. β-DG was used as the loading control. Abbreviations: MHC, myosin heavy chain; β-DG, beta dystroglycan.

## Discussion

The main result of this study is the demonstration that myotubes derived from hiPSC can be used as a common biological resource to study a variety of LGMD. Following terminal differentiation, we reveal here that skMt are sufficiently mature to express the majority of LGMD-causing genes, as well as their related encoded proteins, comparably to immortalized myotubes obtained from muscle biopsies. As a proof of concept study, we further demonstrated that skMt derived from FKRP-deficient hiPSC express the hallmark of LGMDR9 in function of their mutations. Altogether, our results demonstrate that hiPSC represent a valuable unlimited resource for modeling LGMD and assessing *in vitro* efficacy of future LGMD therapies.

While similar in terms of clinical features, LGMD represent a highly heterogeneous group of diseases. Despite the high number of clinical or molecular studies of LGMD subtypes, our current knowledge of the molecular mechanisms underlying these diseases poses many challenges. In this context, the development of a common cellular platform to compare pathological pathways in multiple LGMD would provide a major step forward in their comprehension. So far, hiPSC have been used to explore individual LGMD ^64,66^, but no studies have documented the possibility of using these cells for all types of LGMD. A first study using hiPSC to model one LGMD was reported in 2013 ^49^. In this proof of concept study, the authors showed that LGMD-R2 hiPSC-derived myotubes obtained by stable genomic MyoD1 induction exhibited a defective membrane repair capacity following laser lesion that was rescued by full-length dysferlin gene transfer. More recently, El-Battrawy et al. ^63^ explored ion channel dysfunctions in cardiomyocytes derived from LGMDR9 hiPSC and successfully identified a significant reduction of systolic and diastolic intracellular calcium concentrations in those cells compared to healthy cells. In our study, we explored the possibility of going further by using hiPSC derivatives to model all LGMD genetic subtypes. To do so, we first evaluated the level of expression of all the genes causing LGMD and related proteins in myotubes differentiated from healthy hiPSC. qPCR and immunostaining data revealed that the majority of these genes and proteins were expressed in hiPSC-derived skMt at similar levels to primary myotubes. Beyond the overview of the LGMD that can be studied using hiPSC, this result suggests the use of hiPSC-derived skMt as a platform to improve our understanding of common pathways or phenotypes of several LGMD in a unique cellular model.

As a proof of concept, we then generated and studied hiPSC carrying several mutations of the ambulation, dilated cardiomyopathy, respiratory ^68^ and mental impairment ^69^. From a molecular perspective, FKRP is involved in α-DG glycosylation, as shown by Brockington et al ^70^. While the regulation of α-DG glycosylation appears to be a valuable therapeutic target for the treatment of LGMDR9 ^71^, other studies indicate, on the one hand, that several FKRP mutations do not affect α-DG glycosylation ^72^ and, on the other hand, that there is no correlation between the severity of LGMDR9 and α-DG glycosylation ^73^. Taken together, these studies suggest that the detailed molecular mechanisms underpinning muscular symptoms and pathological pathways in LGMDR9 are still to be elucidated. In our study, we used one hiPSC line provided by the LGMD2I Research Fund and generated three extra hiPSC lines with different mutations. After differentiation into skMt, our results confirmed that the absence of FKRP does not lead to a systematic decrease in α-DG glycosylation in LGMDR9. Only one cell line (FKRP1 with the V300A/A321E mutation) was shown to affect this phenotype, in line with previous studies ^72^.

## Conclusions

In the current study, we demonstrated that skMt derived from hiPSC can be used to model a large panel of LGMD. We showed that almost all genes and proteins involved in LGMD were expressed. We also demonstrated the biological relevance of this model derived from hiPSC through its ability to recapitulate the pathological phenotype identified for LGMDR9. Overall, this study opens up new perspectives on the use of this cellular model to study molecular mechanisms associated with LGMD, but also for the development of high-content phenotypic assays and associated drug screenings.

## Acknowledgments

I-Stem and Genethon are part of the Biotherapies Institute for Rare Diseases, supported by the Association Française contre les Myopathies (AFM-Téléthon). This research was funded by grants from INSERM, the domaine d’intéret majeur (DIM) Biothérapies, université Paris-Saclay, Genopole and the European Commission. The authors thank Marc Peschanski (I-Stem) for helpful discussions, the “technological research team” of I-Stem and Marine Faivre (Genethon) for technical support and the Platform for Immortalization of Human Cells from the Centre of Research in Myology, Institute of Myology, Paris, for providing immortalized muscular cells. In memory of our friend and colleague Jackie Gide.

## Author Contributions

X.N. and I.R. were responsible for the experimental design and project management. C.B. and M.G. performed the characterization of TTN and TTN iPS cells differentiations. C.B. M.G., M.B., E.P. and L.H developed the iPS cells differentiation protocol and carried out molecular analysis of LGMD genes and proteins. M.B. generated, characterized and differentiated FKRP deficient cells. A.B. generated the immortalized cells. C.B., X.N. and I.R. wrote the manuscript.

## Figure legends

**Figure S1.**
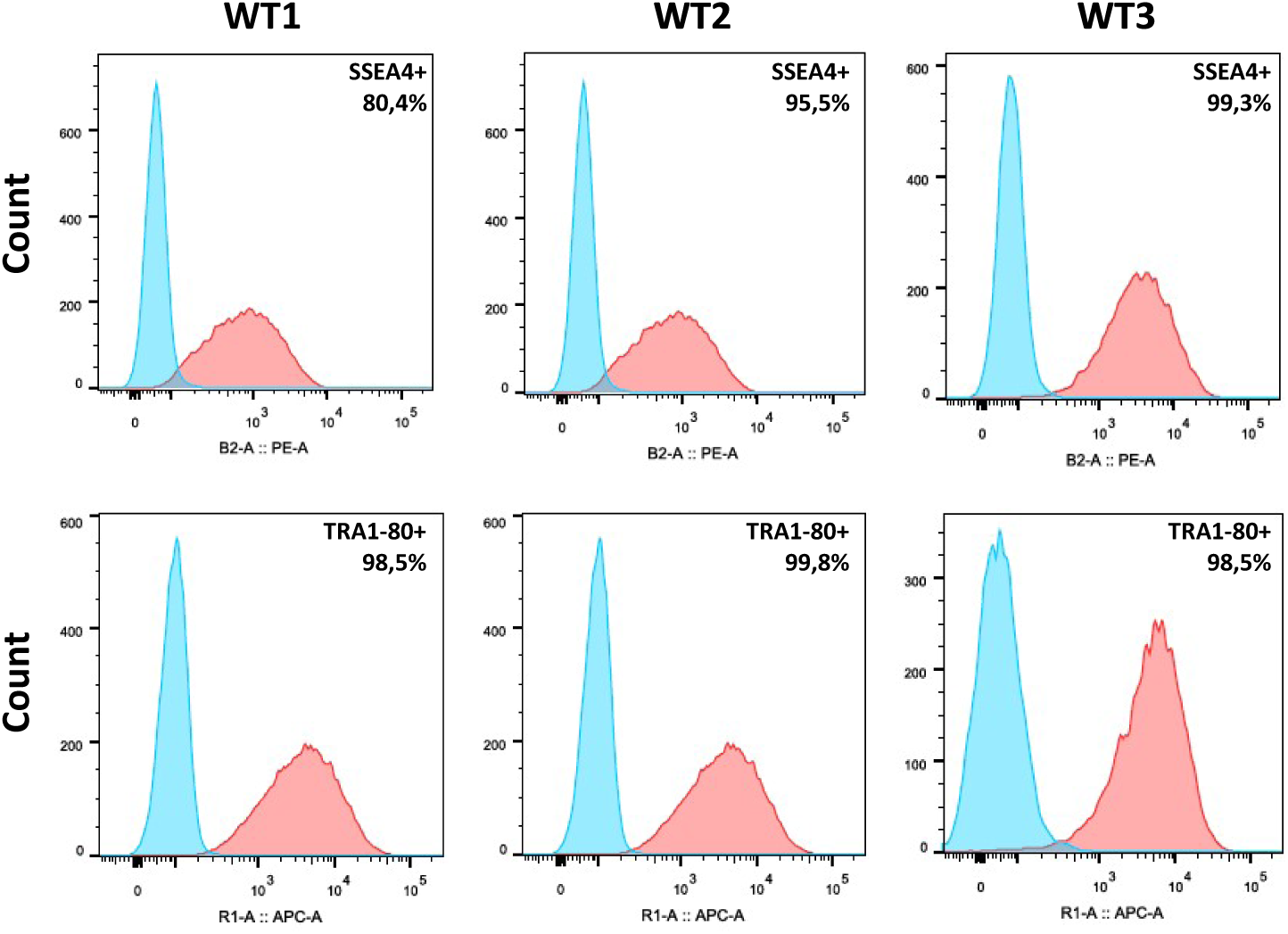
Pluripotency characterization of control hiPSC lines. Flow cytometry analysis of SSEA4 and TRA1-80 in control hiPSC before myogenic differentiation (PE-A+ = SSEA4; APC-A+ = TRA1-80).

**Figure S2.**
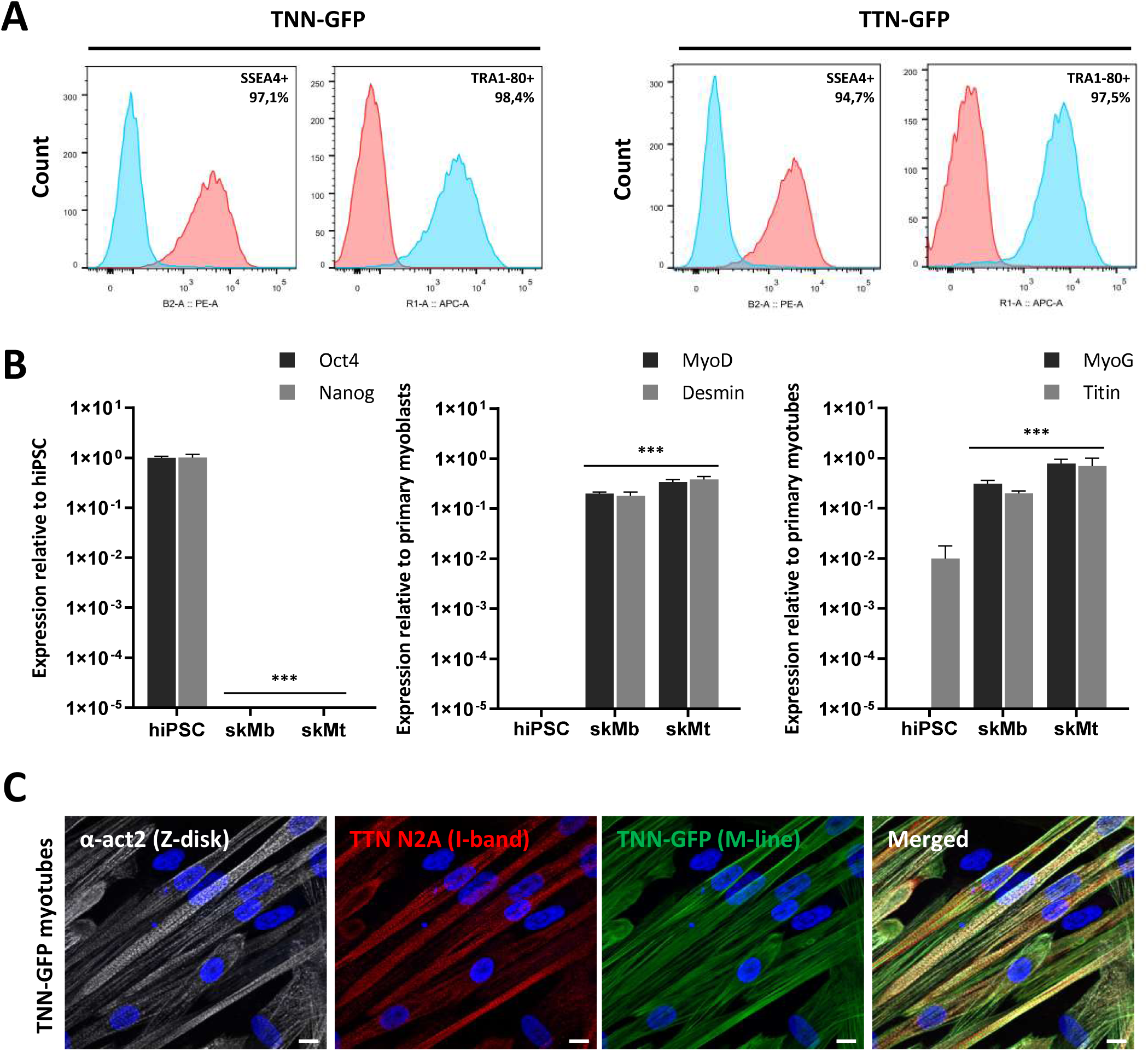
Molecular characterization of TNN- and TTN-eGFP hiPSC and skMC. **(A)** Flow cytometry analysis of SSEA4 and TRA1-80 in TNN- and TTN-eGFP-tagged hiPSC before myogenic differentiation (PE-A+ = SSEA4; APC-A+ = TRA1-80). **(B)** Gene expression analysis of Oct4, Nanog, MyoD, desmin, MyoG and titin in hiPSC, skMb and skMt. Gene expression analyses are normalized to hiPSC, primary myoblasts or primary myotubes. Data are shown as the mean for three independent differentiations of the two tagged cell lines +/- SD. ***p≤0.005 (Student’s t-test). **(C)** Immunostaining analysis by confocal microscopy to visualize sarcomeric structures in TNN-eGFP-tagged skMt at Day 5 of maturation. Nuclei are labelled by Hoechst staining. Scale bar = 10 µm. Abbreviations: MyoG, myogenin; TNN, troponin I; TTN, titin; α-act2, alpha-actinin2.

**Figure S3.**
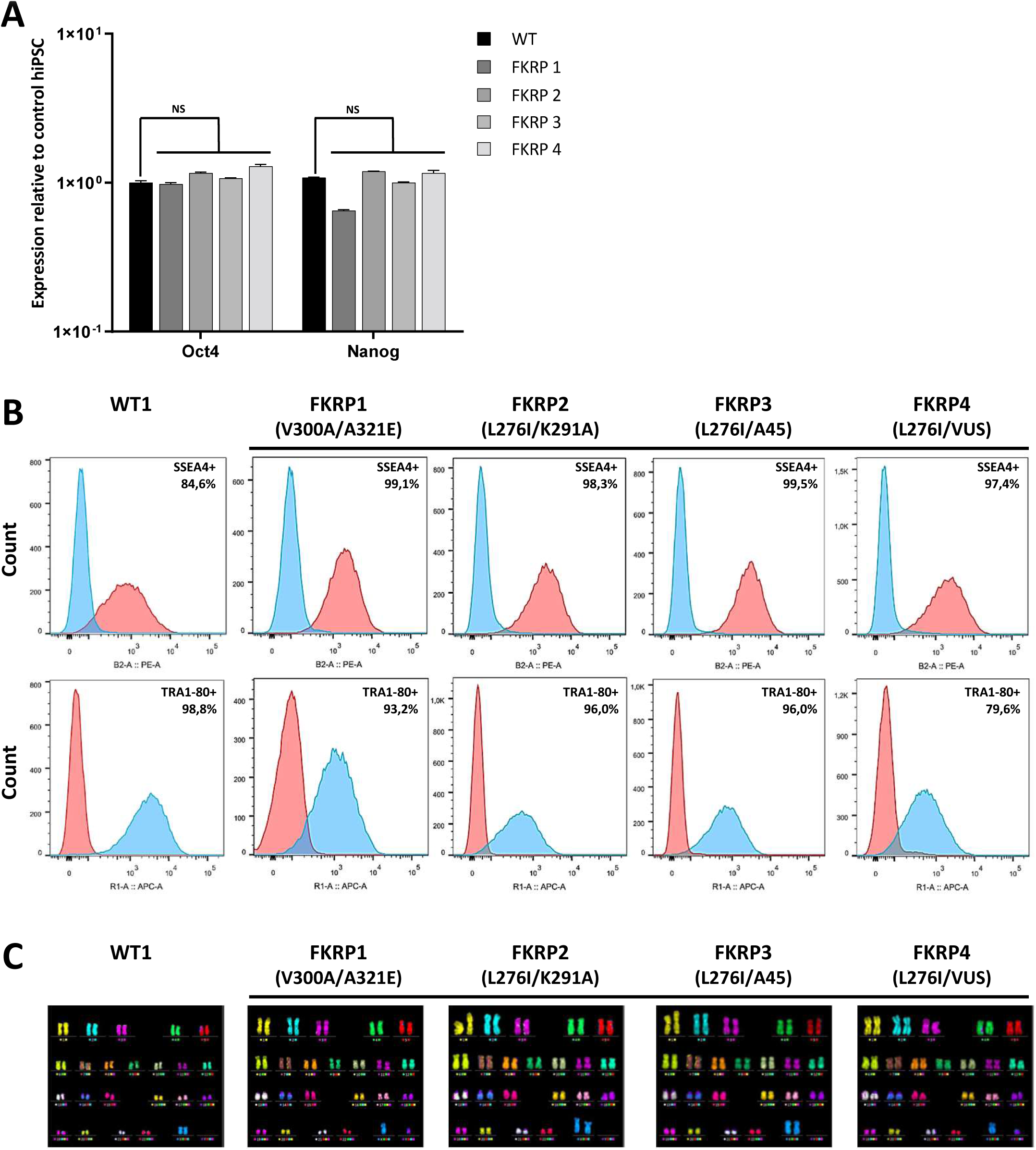
Characterization of control- and LGMDR9-affected hiPSC. **(A)** Gene expression analysis of Oct4 and Nanog in one control and four LGMDR9-affected hiPSC lines. Data are shown as the mean for three independent experiments +/- SD and are normalized to the control cell line. Non-significant p≥0.05 (Student’s t-test). **(B)** Flow cytometry analysis of SSEA4 and TRA1-80 in one control and four LGMDR9-affected hiPSC lines (PE-A+ = SSEA4; APC-A+ = TRA1-80). **(C)** m-FISH analysis of hiPSC lines.

**Figure S4.**
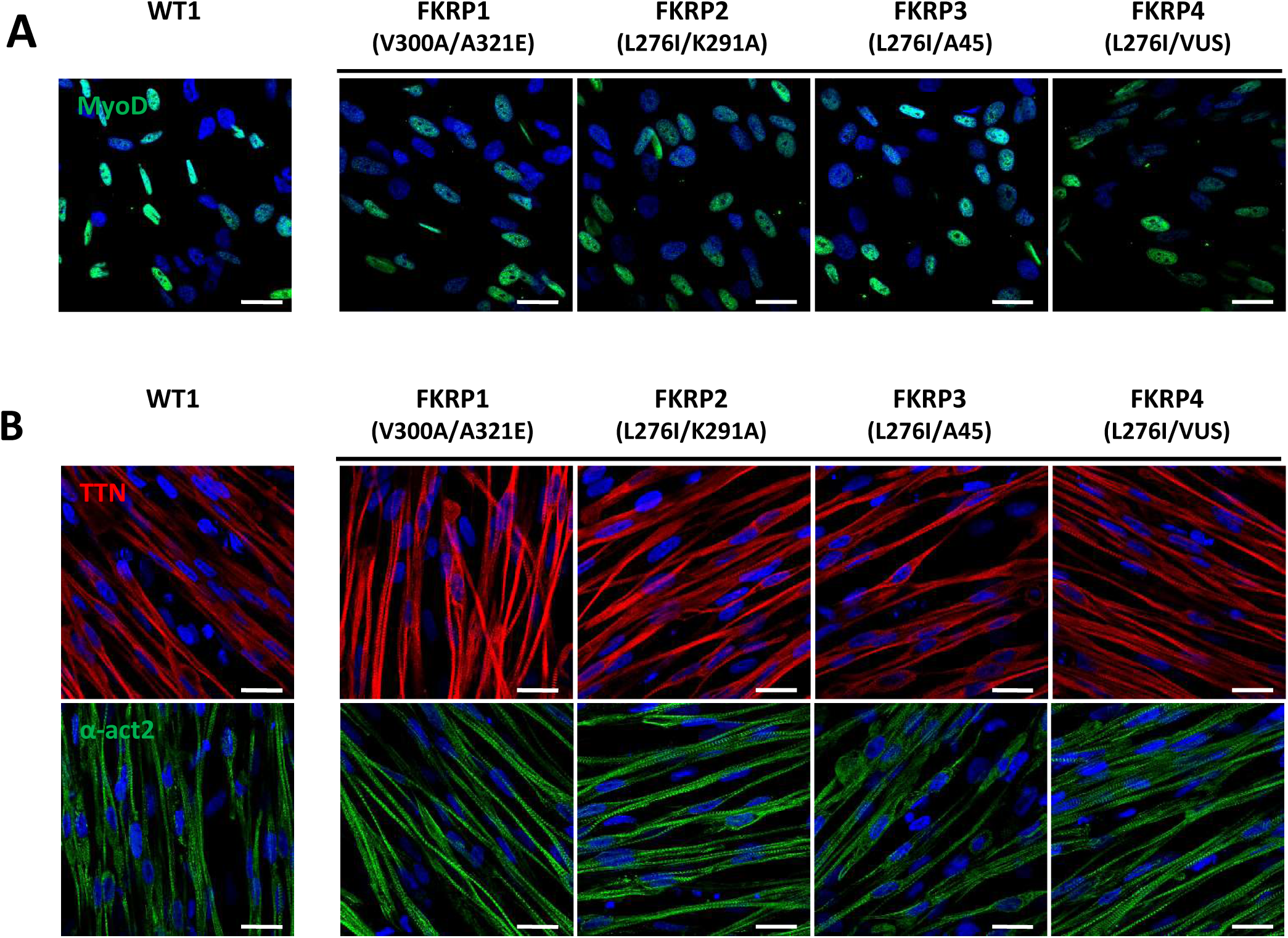
Characterization of FKRP-affected hiPSC differentiation into SkMC. **(A)** Immunostaining analysis of MyoD expression in one control and four LGMDR9-derived skMb at Day 17 of differentiation. **(B)** Immunostaining images of titin and α-actinin in derived skMt at Day 24 of differentiation. Nuclei are labelled by Hoechst staining. Scale bar = 20 µm. Abbreviations: TTN, titin; α-act2, alpha-actinin2.

**Movie S1. Troponin I-eGFP skMt maturation**. Representative movie of TNN-eGFP skMt differentiation from Day 0 to Day 7 after induction of myotube maturation. GFP fluorescence was recorded every 3 hours using the Incucyte® S3 Live-Cell Analysis system (Sartorius) and compiled in a video (4 frames per second).

**Movie S1. Titin-eGFP skMt maturation**. Representative movies of TTN-eGFP skMt differentiation from Day 0 to Day 7 after induction of myotube maturation. GFP fluorescence was recorded every 3 hours using the Incucyte® S3 Live-Cell Analysis system (Sartorius) and compiled in a video (4 frames per second).

**Table S1.** Classification and nomenclature of LGMD.

| New nomenclature | Old nomenclature | Gene | Protein |
| --- | --- | --- | --- |
| LGMD D1 | LGMD 1D | DNAJB6 | DNAJB6, DnaJ heat shock protein family member B6 |
| LGMD D2 | LGMD 1F | TNPO3 | TNPO3, Transportin 3 |
| LGMD D3 | LGMD 1G | HNRNPDL | HNRNPDL, Heterogeneous nuclear ribonucleoprotein D like |
| LGMD D4 / R1 | LGMD 1I / 2A | CAPN3 | Calpain 3 |
| LGMD D5 / R22 | Bethlem myopathy dominant / recessive | COL6A1,<br>COL6A2,<br>COL6A3 | Collagen 6 |
| LGMD R2 | LGMD 2B | DYSF | Dysferlin |
| LGMD R3 | LGMD 2D | SGCA | α-sarcoglycan |
| LGMD R4 | LGMD 2E | SGCB | β-sarcoglycan |
| LGMD R5 | LGMD 2C | SGCG | γ-sarcoglycan |
| LGMD R6 | LGMD 2F | SGCD | δ-sarcoglycan |
| LGMD R7 | LGMD 2G | TCAP | Telethonin |
| LGMD R8 | LGMD 2H | TRIM32 | TRIM32, Tripartite motif containing protein 32 |
| LGMD R9 | LGMD 2I | FKRP | FKRP, Fukutin-related protein |
| LGMD R10 | LGMD 2J | TTN | Titin |
| LGMD R11 | LGMD 2K | POMT1 | POMT1, Protein O-mannosyl-transferase 1 |
| LGMD R12 | LGMD 2L | ANO5 | Anoctamin 5 |
| LGMD R13 | LGMD 2M | FKTN | Fukutin |
| LGMD R14 | LGMD 2N | POMT2 | POMT2, Protein O-mannosyl-transferase 2 |
| LGMD R15 | LGMD 2O | POMGNT1 | POMGNT1, Protein O-linked mannose N-acetylglucosaminyltransferase 1 (beta 1,2-) |
| LGMD R16 | LGMD 2P | DAG1 | Dystroglycan 1 |
| LGMD R17 | LGMD 2Q | PLEC | Plectin |
| LGMD R18 | LGMD 2S | TRAPPC11 | TRAPPC11, Trafficking protein particle complex subunit 11 |
| LGMD R19 | LGMD 2T | GMPPB | GMPPB, GDP-mannose pyrophosphorylase |
| LGMD R20 | LGMD 2U | ISPD | ISPD, 2-C-methyl-D-erythritol 4-phosphate cytidyltransferase |
| LGMD R21 | LGMD 2Z | POGLUT1 | POGLUT1, O-transferase 1 protein |
| LGMD R23 | Laminin α2-related muscular dystrophy | LAMA2 | Laminin α2 |
| LGMD R24 | POMGNT2-related muscular dystrophy | POMGNT2 | POMGNT2, Protein O-Linked Mannose N-Acetylglucosaminyltransferase 2 (Beta 1,4-) |

**Table S2.**
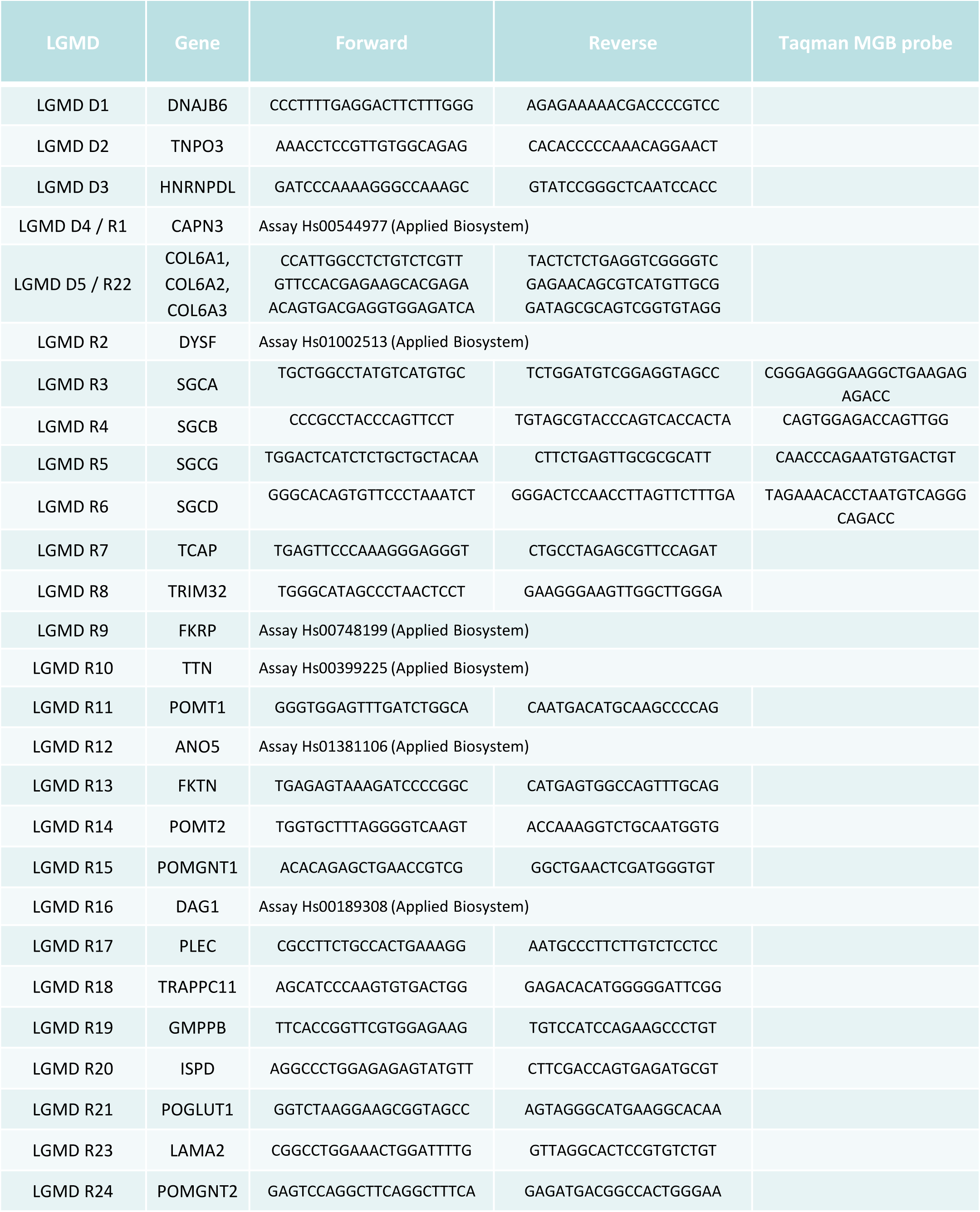
Primers used for the measure of the 27 genes causing LGMD by qPCR.

**Table S3.**
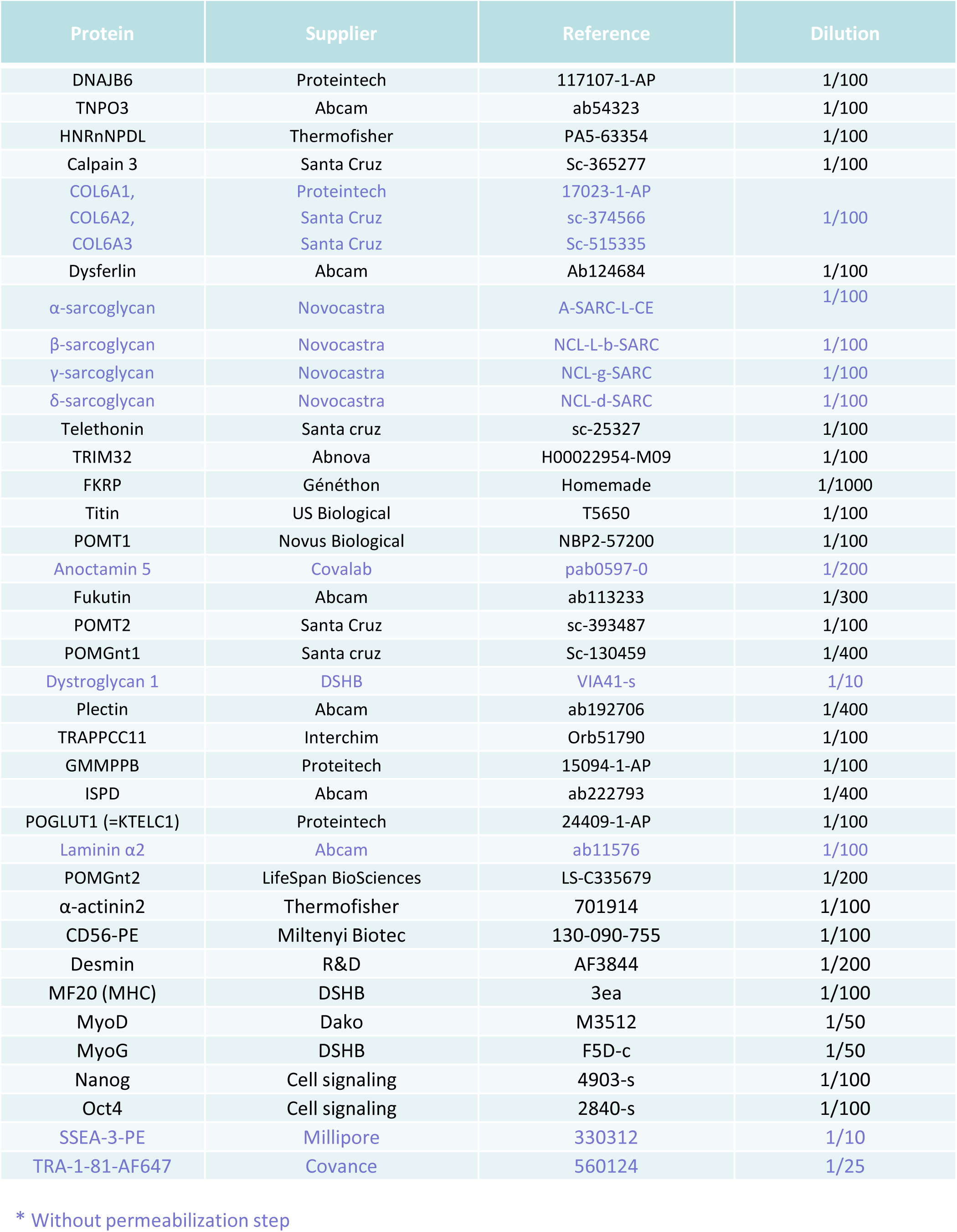
Antibodies used for immunostaining of the 27 proteins causing LGMD.

## Materials and Methods

### Cell lines

Experiments were performed with human immortalized myoblasts from unaffected individuals or with hiPSC from control and LGMDR9-affected patients. The myoblast cell lines were established from human muscular biopsies and obtained from the MyoLine immortalization platform of the Institut de Myologie (Paris, France), with the agreement of the subjects through their signature on an informed consent form and anonymization before immortalization, in line with the EU GDPR regulation. The four cell lines used are the following: C25, AB678, AB1079, AB1167. The WT1 and WT2 hiPSC lines were reprogrammed from healthy IMR-90 lung fibroblasts obtained from the ATCC Cell Lines Biology Collection (Washington, USA) and from GM1869 provided by the Coriell Cell Repository (Camden, USA) ^74,75^. The WT3 iPSC line is a commercial cell line provided by Phenocell (Grasse, France). Wild type eGFP-tagged hiPSC lines, AICS-37 and AICS-48, were provided by the Coriell Cell Repository (Camden, USA): troponinI1 (TNNI1) and titin (TTN) were endogenously tagged with mEGFP at the C-terminus site using CRISPR-Cas9 technology. The FKRP1 iPSC line was reprogrammed by Phenocell (Grasse, France) from LGMDR9 fibroblasts (heterozygous FKRP V300A/A321E) provided by Genethon’s Cell Bank (Evry, France). FKRP2 and FKRP3 iPSC lines were also reprogrammed by Phenocell (Grasse, France) from LGMDR9 fibroblasts (heterozygous L276I/K291A and L276I/A45, respectively) provided by the Coriell Institute (Camden, USA). The FKRP4 iPSC line (heterozygous L276I/VUS) was provided by the LGMD2I Research Fund.

### Cell culture and differentiation

Immortalized myoblasts were cultivated in a growth medium: 1 volume of 199 medium (Invitrogen) for 4 volumes of DMEM (Invitrogen), supplemented with 20% FCS (Sigma), 25 µg/ml fetuin (Life Technologies), 5 ng/ml hEGF (Life Technologies), 0.5 ng/ml bFGF (Life Technologies), 0.2 µg/ml dexamethasone (Sigma) and 5 µg/ml insulin (Sigma). Cells were seeded on 0.1% gelatin and maintained in a humidified atmosphere of 5% CO2 at 37°C. Differentiation was induced at confluence by replacing the growth medium with DMEM supplemented with 10 μg/ml insulin (Sigma) for 7 days. Control and LGMDR9 hiPSC lines were maintained and expanded using a single cell method on feeder-free coated dishes (Matrigel, Corning) with iPS-Brew XF medium (Macs, Milteny Biotech). hiPSC were differentiated into skMC following a protocol developed by Geneabiocell ® (Caron et al., 2016). Briefly, hiPSC were seeded in collagen I-coated plates (Biocoat, DB Biosciences) and maintained for 10 days in skeletal muscle induction medium (SKM01, AMSBIO) with a passage at Day 7. Cells were then dissociated with 0.05% trypsin and seeded once again onto collagen I-coated plates for 7 days in skeletal myoblast medium (SKM02, AMSBIO) until freezing. skMb were thawed on collagen I-coated glass slides in skeletal myoblast medium and incubated at confluence with skeletal muscle differentiation medium (SKM03, AMSBIO) for 5 to 7 supplementary days. hiPSC, skMb and skMt were analyzed at Days 0, 17 and 24 of differentiation, respectively.

### Flow cytometry

A single-cell suspension of hiPSC was collected after chemical dissociation with accutase (Invitrogen), centrifuged at 900 rpm for 5 min and resuspended in 2% FBS in cold PBS. Cells were stained with fluorescent dye-conjugated antibodies for 30 min on ice and protected from light. Antibodies: APC-conjugated TRA1-80 (Millipore) and PE-conjugated SSEA4 (Millipore). Cells were washed in cold PBS before being sorted by a MACSquant analyzer (MiltenyiBiotec). Data were analyzed with FlowJo Software (Tree Star).

### Multiplex fluorescence in situ hybridization (m-FISH) karyotype analysis

Cells were blocked in metaphase with colchicine (Eurobio) for 90 min, warmed with a hypotonic solution and fixed with a Carnoy fixative. An M-FISH 24Xcite probe (MetaSystems) and ProLong Gold Antifade Mountant with DAPI (Thermo Fisher Scientific) were used for m-FISH staining. 70 metaphases were acquired with Metafer MetaSystems software coupled to an AxioImager Z2 (Zeiss) microscope equipped with a camera cool cube and 10X and 63X objectives. Images were analyzed with Isis software (MetaSystems).

### Immunofluorescence analysis

Cells at Days 0, 17 or 24 of skMC differentiation were fixed with 4% paraformaldehyde for 10 min at room temperature. For two antibodies (FKTN and TCAP), cells were pre-fixed and then fixed in cold methanol for 5 min at room temperature. After 3 washes in phosphate-buffered saline (PBS), cells were permeabilized, or not, with 0.5% Triton X-100 for 5 min and blocked in PBS solution supplemented with 1% bovine serum albumin (BSA, Sigma) for 1 hour at room temperature. Cells were stained for specific markers overnight at 4°C using the antibodies listed in Table S3. After 3 washes in PBS, staining was revealed by appropriate Alexa Fluor secondary antibodies (ThermoFisher Scientific Life Sciences; 1:1000) in the dark for 1 hour at room temperature and nuclei were visualized with Hoechst 33342 (Invitrogen; 1:2000). Cell imaging was carried out with a Zen Black software-associated LSM-800 confocal microscope (Zeiss) with a 20x or 63X objective.

### Time lapse imaging of skMt differentiation

Time-lapses of skMt were recorded as previously described, from Day 0 to Day 5 of differentiation. Imaging of endogenous GFP-tagged titin and troponin signaling was carried out with an Incucyte® S3 Live-Cell Analysis system (Sartorius) or a LSM-800 confocal microscope (Zeiss) with a 20x objective, using Zen Black software. Differentiation efficiency was measured as the percentage of GFP-tagged cells at each time point using ImageJ software and normalized to the total number of skMt. An algorithm was used to determine the percentage of cells when at least 50% of the area of an identified nucleus was covered by GFP staining. Differentiation efficiency was calculated from 5 random fields comprising at least 300 cells. The results are expressed as mean values +/- S.E.M. *** p<0.005 (Student’s t-test).

### Quantitative PCR

Total RNAs were isolated using the RNeasy Mini extraction kit (Qiagen) according to the manufacturer’s instructions. A DNase I digestion was performed to degrade DNA in the sample. RNA levels and quality were checked using the NanoDrop technology. A total of 500 ng of RNA was used for reverse transcription using the SuperScript III reverse transcription kit (Invitrogen). Quantitative polymerase chain reaction (qPCR) analysis was performed using a QuantStudio 12CK Flex real-time PCR system (Applied biosystem) and Luminaris Probe qPCR Master Mix (Thermo Scientific), following the manufacturers’ instructions. Gene expression analysis was performed using the TaqMan gene expression Master Mix (Roche), following the manufacturer’s protocol. Quantification of gene expression was based on the DeltaCt method and normalized to 18S expression (Assay HS_099999). The primers and probe sequences that were designed are listed in **Table S2**. The sequences of the remaining genes are commercially available (Applied biosystem):

DYSF (Hs01002513), FKRP (Hs00748199), TTN (Hs00399225), ANO5 (Hs01381106) and DAG1 (Hs00189308).

### Western immunoblotting

Whole-cell lysate of control and LGMDR9-skMt were collected after seven days of differentiation. Proteins were extracted with RIPA lysis buffer (Thermo Scientific) supplemented with 1X Protease Inhibitor-Complete ULTRA tablets mini (Roche) and 1X benzonase nuclease HC (Millipore) for 1⍰h at 4⍰°C. Protein concentration was measured using the Pierce BCA Protein Assay Kit (ThermoScientific) and the absorbance at 562 nm was evaluated using a CLARIOstar® microplate reader (BMG Labtech). For α-DG protein detection, a total of 130Cμg of protein was loaded and run on Mini-PROTEAN 4-12% bis-tris protein gels (BioRad) and then transferred to nitrocellulose membranes with a Trans-Blot Turbo Transfert system (Biorad), following the manufacturer’s instructions. Membranes were then blocked in Odyssey blocking buffer (Li-Cor) for 1 hour at room temperature. Incubation with primary antibodies diluted in Odyssey blocking buffer was carried out at 4°C from 2 hours to overnight for the mouse anti-α-DG-IIH6 1:50 (DSHB), the mouse anti-α-DG-IIH6 1:1000 (Merck Millipore) and the mouse anti-β-DG 1:200 (DSHB). Washing was carried out 3 times for 10 minutes at room temperature with TBS + 0.1% Tween20 and the membranes were incubated with a donkey anti-mouse antibody IRDye-800CW 1:5000 (Li-Cor) in blocking buffer. Washing was carried out and proteins were detected by fluorescence (Odyssey, Li-Cor), following the manufacturer’s instructions.

### Statistical analysis

Data are presented as means ±⍰SD unless otherwise specified. Statistical analysis was performed using the Student’s t test. *P* values ≤ 0.05 (*), 0.01 (**) and 0.005 (***) were considered significant.

